# Phytosphingosine induces systemic acquired resistance through activation of sphingosine kinase

**DOI:** 10.1101/2020.05.08.084657

**Authors:** So Yeon Seo, Yu Jung Kim, Myung Hee Nam, Ky Young Park

## Abstract

Phytosphingosine (PHS) is a naturally occurring bioactive sphingolipid molecule. Intermediates such as sphingolipid long-chain bases (LCBs) in sphingolipid biosynthesis have been shown to have important roles as signaling molecules. In this study, exogenous addition of PHS caused rapid induction of transcripts responsible for transient synthesis of LCBs, reactive oxygen species, and ethylene. These events were followed by the induction of sphingolipid kinase (SphK), which metabolized PHS to phytosphingosine-1-phosphate in an biphasic manner. PHS alleviated not only pathogen-induced cell damage but also reduced the growth of virulent pathogens in the entire upper part of the PHS-treated plant stem during the necrotic stage after inoculation, suggesting the development of systemic acquired resistance (SAR) and plant immunity. Moreover, PHS treatment up-regulated the transcription and activity of SphK, accompanied by prominent increases in the transcription levels of serine palmitoyltransferase (*LCB1* and *LCB2*) for *de novo* synthesis of sphingolipids, as well as ROS-detoxifying enzymes and PR proteins at 48 h after virulent pathogen infection. The impairment of ROS production at this time is more beneficial for the activation of SphK and inhibition of pathogenicity during the necrotic stage of hemibiotrophic infection, indicating that necrotic cell death at the late stage is regulated by ROS-independent SphK. Phosphorylated LCBs significantly reduced pathogen-induced cell damage. These observations suggest that selective channeling of sphingolipids into phosphorylated forms in a time-dependent manner has a pro-survival effect by promoting SAR in plant immunity.

**One Sentence Summary:** Selective gene expression in sphingolipid biosynthesis and channeling into their phosphorylated forms are significant determinants of their roles as pro-survival signaling molecules.

## Introduction

Sphingolipids are essential structural components of the cellular membrane system, which includes the plasma membrane, tonoplast, and other endomembranes of plant cells that provide mechanical stability (Markham et al., 2006). It is estimated that sphingolipids constitute up to 30% of the tonoplast and plasma membrane lipids in plants (Lynch and Dunn, 2004). Further, sphingolipids are reported to participate as bioactive molecules in the regulation of intracellular processes including cell proliferation, differentiation, development, apoptosis, angiogenesis, and immunity in all eukaryotic and some prokaryotic cells (Aguilera-Romero et al., 2013). Moreover, sphingolipid metabolites have recently drawn attention as second messengers for stomatal closure, programmed cell death (PCD), and defense against pathogen attack in response to abiotic and biotic stresses in plants (Markham et al., 2013; Magnin-Robert et al., 2015; Glenz et al., 2019). Although the physiological roles of sphingolipid have been extensively studied in animals (Mashima et al., 2020), it remains unclear what mechanism is responsible for their physiological action in plants.i

Long-chain bases (LCBs) are unique building blocks in all sphingolipids including ceramide, sphingomyelin, dihydrosphingosine, and phytosphingosine. Sphingolipid biosynthesis is initiated in the endoplasmic reticulum (ER) by condensation of serine with palmitoyl-coenzyme A catalyzed by serine palmitoyltransferase (SPT), resulting in the production of the LCB 3-ketodihydrosphinganine, which is then reduced to dihydrosphingosine (d18:0) (Chao et al., 2011). Dihydrosphingosine is further modified to phytosphingosine (t18:0) (PHS) by the introduction of a double bond into the LCB in plants and fungi (Berkey et al., 2012), as well as to sphingosine, a free form of long-chain sphingoid base, in mammals (Mashima et al., 2020). The highest proportion of PHS, a trihydroxylated LCB, has been reported in the leaves of tobacco (*Nicotiana tabacum* and *Nicotiana benthamiana*) with large proportions of d18:2 and t18:1 (Cacas et al., 2012). Therefore, the composition of sphingolipids in plants is notably different from those in animals and fungi, in which the major LCBs are t18:1 and d18:2.

One very important sphingolipid metabolite is sphingosine-1-phosphate (S1P), which is formed from sphingosine by sphingosine kinase (SphK). The activity of S1P has been described in a wide spectrum of organisms, ranging from *Arabidopsis thaliana* through *Saccharomyces cerevisiae* to *Homo sapiens* (Bourquin et al., 2010). S1P has recently received attention due to its regulation of many biological processes such as cell growth and survival, proliferation, differentiation, and immune function in eukaryotes (Spiegel and Milstien, 2003; Proia and Hla, 2015). S1P functions as an intracellular second messenger and a ligand for G-protein-coupled cell-surface receptors belonging to the lysophospholipid receptor family in mammals, which functions that have been implicated in cell growth and inhibition of apoptosis (Spiegel and Milstien, 2003). It was recently reported that S1P signaling via sphingosine 1-phosphate receptor-1 (S1PR1), a cell-surface receptor, can enhance tumorigenesis and stimulate growth, expansion, angiogenesis, metastasis, and survival of cancer cells (Cartier et al., 2020). Therefore, potential uses of S1P signaling modulators as pharmaceutical and therapeutic targets in cancer therapy have been suggested.

The evidence for S1P as a signaling molecule in mammals has been extended to plants. S1P is involved in abscisic acid (ABA)-mediated guard cell signal transduction in response to chilling (Dutilleul et al., 2012) in *Arabidopsis*. Phytosphingosine-1-phosphate (PHS1P), similar to S1P, is active in stress signaling, which is mediated by the G-protein α-subunit in *Arabidopsis* (Coursol et al., 2005). More recent studies have observed that they participate as signaling molecules in the defense pathways of the hypersensitive response (HR) (Glenz et al., 2019).

The final step in the sphingolipid degradative pathway is mediated by S1P lyase (SPL), which irreversibly converts S1P to hexadecenal and phosphoethanoamine (Bandhuvula and Saba, 2007). It has been suggested that SPL silences the sirens from the immune system by removing the available S1P signaling pools. However, there is the reversible dephosphorylation of S1P back to sphingosine, which is mediated by S1P phosphatase (SPP) (Zhang et al., 2012). Both SPL and SPP regulate LCB/LCBP homeostasis. Recent studies have suggested new opportunities for the use of S1P-metabolizing enzymes as antiviral drugs by targeting the host enzymes (Wolf et al., 2019). Similar findings that modification of sphingolipid metabolite content can affect plant defense responses have been reported in *Arabidopsis* (Magnin-Robert et al., 2015).

PHS is a major LCB in some plants and is involved in cell signaling. In a recent study, sphingolipidomic profiling revealed that PHS accumulates as early as 1 h after pathogen inoculation with virulent and avirulent strains of *Pseudomonas syringae* pv. *tomato* in *Arabidopsis* leaves (Peer et al., 2010). We previously reported that PHS levels rapidly increased in susceptible tobacco (*Nicotiana tabacum* L. cv Wisconsin 38) plants at 1 h and 48 h after shoot inoculation with the hemibiotrophic pathogen *Phytophthora parasitica* var. *nicotianae* (*Ppn*), which was determined by performing ultraperformance liquid chromatography-quadrupole-time of flight/mass spectrometry (Cho et al., 2012). Herein, we tried to establish a pathophysiological link between pathogen infection and sphingolipid metabolism in tobacco plants. Further, we analyzed the significance of sphingolipid metabolites such as PHS, PHS1P, and S1P during plant-pathogen interactions and their contribution to plant defense.

## Results

### PHS-induced rapid signaling responses

Although the physiological roles of sphingolipids are not fully described, recent studies have indicated that they have crucial roles in the induction of apoptotic-like cell death in plants (Berkey et al., 2012). Initially, we determined the rate of cell damage in PHS-treated tobacco plant stems with 5 leaves, followed by photography after staining with lactophenol trypan blue. PHS was treated for 12 h at concentrations of 1 μM to 10 μM, and the results showed that PHS treatment slightly induced cell death beginning at 1 μM PHS (Fig. 1A). Even though the tobacco leaves were not severely damaged at 10 μM PHS, as the concentration of PHS increased, the degree of damage to plant leaves slightly increased.

**Figure 1.**
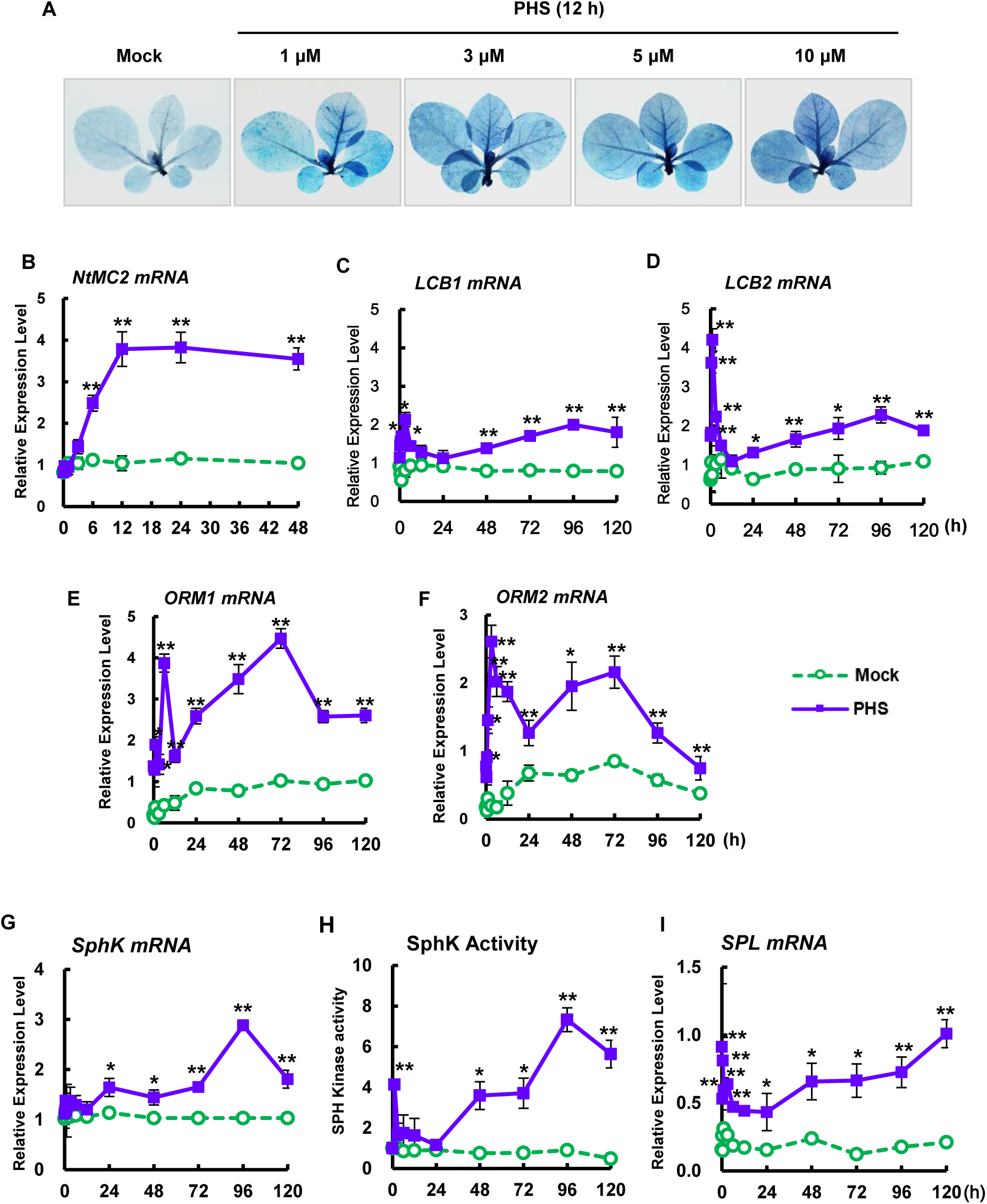
Effect of PHS treatment on cell damage and sphingolipid metabolism in tobacco leaves. A, Mature tobacco leaves were treated with different concentrations of PHS for 12 h, and after the indicated time necrotic areas were stained with trypan blue and then imaged with a digital camera. B, Transcription levels of tobacco *NtMC2* gene in tobacco plants after PHS treatment for 48 h. Results of real-time qRT-PCR analysis of *NtMC2* transcription after 1 μM PHS treatment using total RNAs from tobacco leaves. C-F, Transcription levels of *LCB1* (C) and *LCB2* (D) of serine palmitoyltransferase and *ORM1* (E) and *ORM2* (F) of orosomucoid in tobacco leaves after 1 μM PHS treatment. G-H, Transcription level (G) and activity (H) of sphingosine kinase in tobacco leaves after 1 μM PHS treatment. I, Transcription levels of *SPL* of sphingosine-1-phoshpate lyase in tobacco leaves after 1 μM PHS treatment. Transcription levels are expressed relative to the reference gene *β-actin* after real-time qRT-PCR. Relative mRNA expression levels are expressed as means ± SD. One asterisk (P < 0.05) or two asterisks (P < 0.01) indicate a significant difference between mock- and PHS-treated cases at the same indicated time.

Therefore, we investigated the expression of genes related to cell damage after treatment with 1 μM PHS, which produced relatively weak cell damage. We first determined the effects of PHS treatment on the expression of the metacaspase type II gene (*MC2*), which mediates biotic and abiotic stress-induced PCD (Watanabe and Lam, 2011). Following PHS treatment, the transcription level of *NtMC2* gradually increased to a maximum level at 12 h (Fig. 1B), after which it decreased somewhat until 48 h. Therefore, the absence of a further increase in *NtMC2* transcription after 12 h indicates that cell damage did not progress into a severe state, which is in accordance with the PHS-induced cell death pattern (Fig. 1A). These results indicate that a rapid up-regulation of *NtMC2* expression is responsible for the PHS-mediated HRs at an early stage.

We next determined whether alteration of sphingolipid metabolism occurs after PHS treatment. LCB molecules, which are unique components of sphingolipids, are formed by condensation of serine and palmitoyl-coenzyme A (Dietrich et al., 2008). This reaction is the first distinctive step in sphingolipid biosynthesis, which is catalyzed by the pyridoxal phosphate-dependent enzyme SPT (EC 2.3.1.50). SPT, an ER-associated heterodimeric protein consisting of LCB1 and LCB2 subunits, is thought to be a rate-limiting enzyme in the sphingolipid biosynthetic pathway (Dietrich et al., 2008). It was recently reported that PHS is sufficient to induce ER stress surveillance phenotypes in budding yeast, *S. cerevisiae*, which in turn elevates the PHS level, suggesting the presence of feedback activation in pathogen-induced sphingolipid biosynthesis.

Gene expression levels of LCB1 and LCB2 subunits were determined in order to elucidate the effects of PHS in sphingolipid biosynthetic pathways (Fig. 1, C and D). Although transcriptions of *LCB1* and *LCB2* were unaltered in mock-treated control plants, transcription levels of both genes were rapidly up-regulated from 15 min after PHS treatment, with higher expression of *LCB2* than *LCB1*. PHS-induced activation had the strongest effects on LCB1 at 3 h and on LCB2 at 1 h. The amounts of both transcripts and their abundance patterns from 24 h to 120 h were similar, with second peaks occurring at 96 h after PHS treatment. The similar biphasic pattern was also detected in the transcript changes of *ORM1* and *ORM2* of tobacco orosomucoid (Fig. 1, E and F), which mediate SPT oligomerization and its subcellular localization, but, may directly or indirectly inhibit its activity (Han et al., 2020). These results indicate that *de novo* synthesis of sphingolipids occurs through feedback activation induced by exogenously added PHS. Our results support the previous report that PHS-induced co-activation of *LCB1* and *LCB2* is responsible for the biosynthesis of bioactive sphingolipids that regulate many cellular responses (Proia and Hla, 2015).

We next investigated the effects of PHS treatment on the transcription and activity of SphK, which catalyzes PHS to PHS1P, which is abundant in fungi and plants (Dutilleul et al., 2012). PSH1P has a fundamental role as an intracellular signaling molecule in development of and stress responses in eukaryotic cells (Piña et al., 2018). Although the transcription level of *SphK* was immediately increased after PHS treatment, it returned to the basal level after 12 h (Fig. 1G). The amount of *SphK* transcripts began to increase again after 24 h of PHS treatment, reaching a maximum at 96 h and then decreasing. However, no change was observed for the entire period following mock treatment.

The level of SphK activity responded to PHS treatment in a biphasic manner (Fig. 1H). SphK activity began to increase rapidly and peaked at 1 h, after which it returned to the basal level. However, it increased again after 24 h, and reached a peak at 96 h, similar to the maximum pattern level of the *SphK* transcript (Fig. 1H). These results indicate that PHS-induced SphK is transcriptionally regulated. It is suggested that exogenously added PHS can, at a later stage, be metabolized significantly to PHS1P, which is known to be involved in intracellular signaling (Coursol et al., 2005).

The final step in sphingolipid metabolism is mediated by SPL, which degrades long-chain base-1-phosphates (LCBPs) such as PHS1P and is the only reported route for the destruction of sphingolipids (Bandhuvula and Saba, 2007). The amount of *SPL* mRNA induced after PHS treatment was biphasic (Fig. 1I). The amount of *SPL* mRNA rapidly increased to a peak at 0.5 h after PHS treatment; after which, it lowered to basal levels before gradually increasing up to 120 h.

### NADPH oxidase-dependent transient ROS accumulation at an early stage after PHS treatment

It has been recognized that ROS generation and signaling can activate HR-related localized cell death in plants (Zurbriggen et al., 2010). PHS is considered a powerful contributor to oxidative damage in eukaryotic cells. To investigate ROS generation in tobacco leaves following treatment with 1 μM PHS, we histochemically monitored the levels of two important ROS, superoxide anion and hydrogen peroxide, by using 3,3’-diaminobenzidine (DAB) and nitrotetrazolium blue chloride (NBT) as chromogenic substrates, respectively, in tobacco leaves for up to 48 h. NBT reacts with O_2_^−^ to form a dark blue insoluble formazan compound, and DAB is oxidized by hydrogen to produce a reddish-brown precipitate (Kumar et al., 2014). Both ROS were significantly detected from 15 min after PHS treatment and abundance peaked at 1 h, suggesting the ROS were rapidly and transiently generated (Fig. 2, A and B). Surprisingly, only very low levels of both ROS were produced after 3 h (Supplemental Fig. S1). These results imply that PHS-induced ROS generation is only HR-relevant at an early stage.

**Figure 2.**
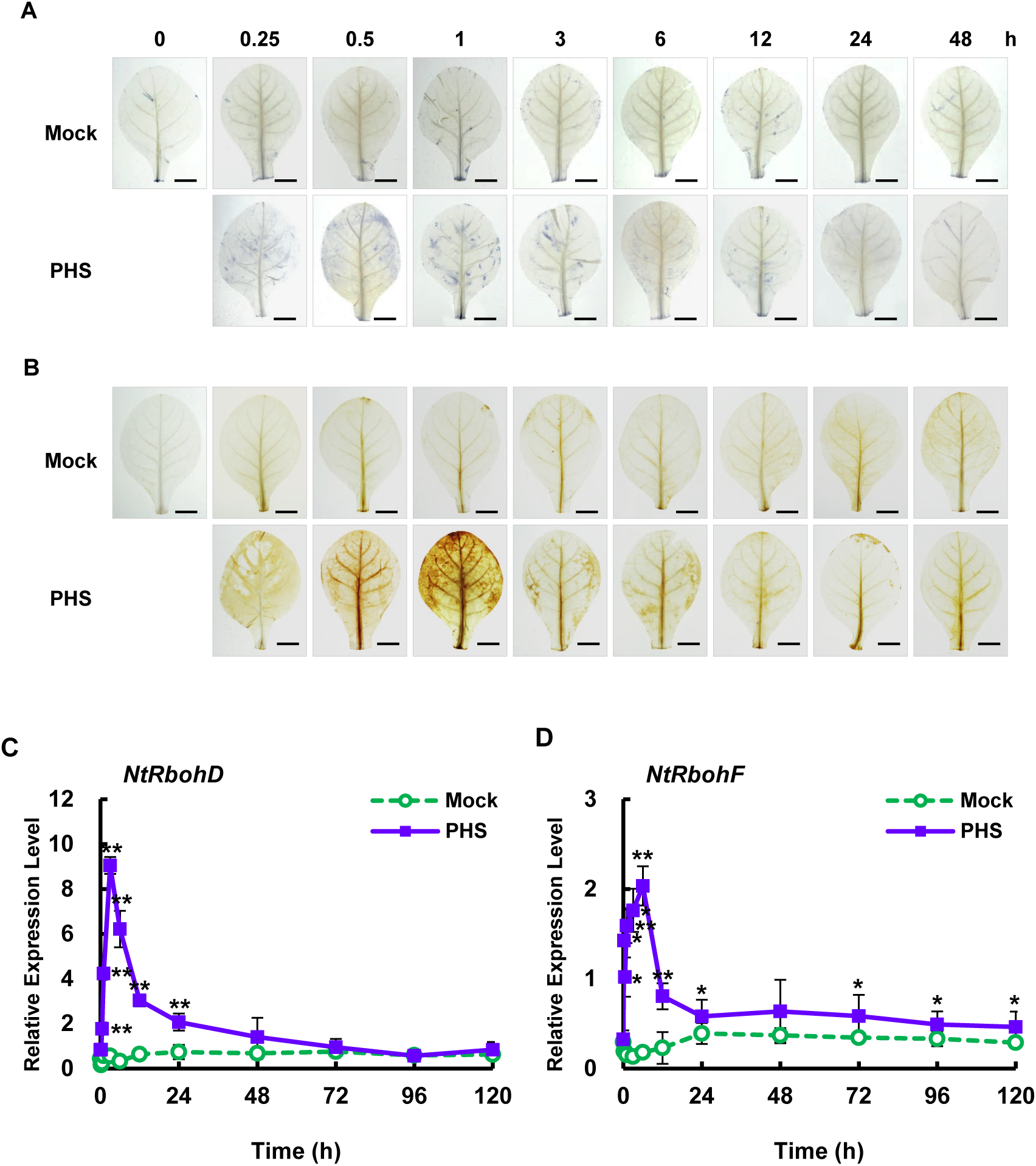
Effects of PHS treatment on ROS accumulation and *NADPH oxidase* gene (*NtRbohD* and *NtRbohF*) transcription in tobacco leaves. A-B, Histochemical analysis of intracellular ROS accumulation in PHS-treated tobacco leaves. After mature tobacco leaves were treated with 1 μM PHS for 48 h, superoxide anion level was determined by NBT staining (A), and H_2_O_2_ level was detected by DAB staining (B). Staining images of leaves were photographed by a digital camera. C-D, Relative mRNA levels of *NtRbohD* and *NtRbohF* genes in mature tobacco leaves treated with 1 μM PHS. Transcription levels of *NtRbohD* (C) or *NtRbohF* (D) are expressed as means ± SD. Transcription levels are expressed relative to the reference gene *β-actin* after qRT-PCR. An asterisk indicates a significant difference between mock- and PHS-treated cases (one asterisk (P < 0.05) or two asterisks at the same time point (P < 0.01)).

In aerobic organisms, ROS are produced during normal cellular metabolism as by-products of metabolic pathways and electron flow in both mitochondria and chloroplasts. The neutrophil nicotinamide adenine dinucleotide phosphate (NADPH) oxidase, RboH, is a transmembrane protein that generates superoxide radicals in plant cells. Its isoforms, *NtRbohD* and *NtRbohF*, are expressed throughout tobacco plants and produce superoxide anions, which are unable to permeate cell membranes under ambient pH conditions due to the presence of a negative charge.

To provide further insight into ROS accumulation in PHS-treated tobacco leaves, we examined gene expression profiles of tobacco plants in the context of ROS production following PHS treatment. To determine the expression of *NtRboh* genes in PHS-induced ROS generation, we measured the transcription levels of two *NtRboh* genes in tobacco leaves using real-time qRT-PCR analysis. Transcription of *NtRbohD* and *NtRbohF* occurred in a time-dependent manner; more specifically, *NtRbohD* and *NtRbohF* transcription showed monophasic kinetics upon PHS treatment, displaying early transient peaks at 3 h and 6 h, respectively (Fig. 2, C and D). The transcription level of *NtRbohD* increased by about 7.4-fold in the PHS-treated tobacco plants compared to that in mock-treated plants at 1 h after PHS treatment. Soon thereafter, *NtRbohD* transcription gradually decreased, returning to almost basal level after 48 h. The relative transcription level of *NtRbohD* at 1 h normalized to the expression level of *β-actin* was about four times higher than that of *NtRbohF* at 1 h, suggesting that *NtRbohD* is more responsive than *NtRbohF* during PHS-induced ROS generation.

### PHS-induced ethylene production is dependent on *NtACS2* and *NtACS4* expression

Ethylene is implicated as a virulence factor of pathogens as well as a signaling molecule in disease resistance (Wi et al., 2012). Pathogen infection was shown to induce typical responses, including biphasic production of ROS and ethylene, in which synergism between ROS and ethylene constitutes an important regulatory mechanism in tobacco leaves (Wi et al., 2012). Therefore, we propose that ethylene production could be a central component of a self-amplifying loop wherein transient biphasic ROS bursts have a critical regulatory role.

To further determine whether or not ethylene functions as a physiological amplifier in PHS-induced cell death, we measured ethylene production after PHS treatment. Monophasic ethylene production was observed in tobacco leaves after PHS treatment (Fig. 3A), which was in accordance with the observation of monophasic ROS production (Fig. 2, A and B). Ethylene production rapidly increased at 30 min, reached a transient peak at about 1 h, and declined thereafter. After 30 h of PHS treatment, ethylene production had completely returned to the basal level. Taken together, the results show that PHS-induced ROS generation is followed by PHS-induced ethylene production, suggesting that ROS generation functions upstream of ethylene generation in response to PHS treatment.

**Figure 3.**
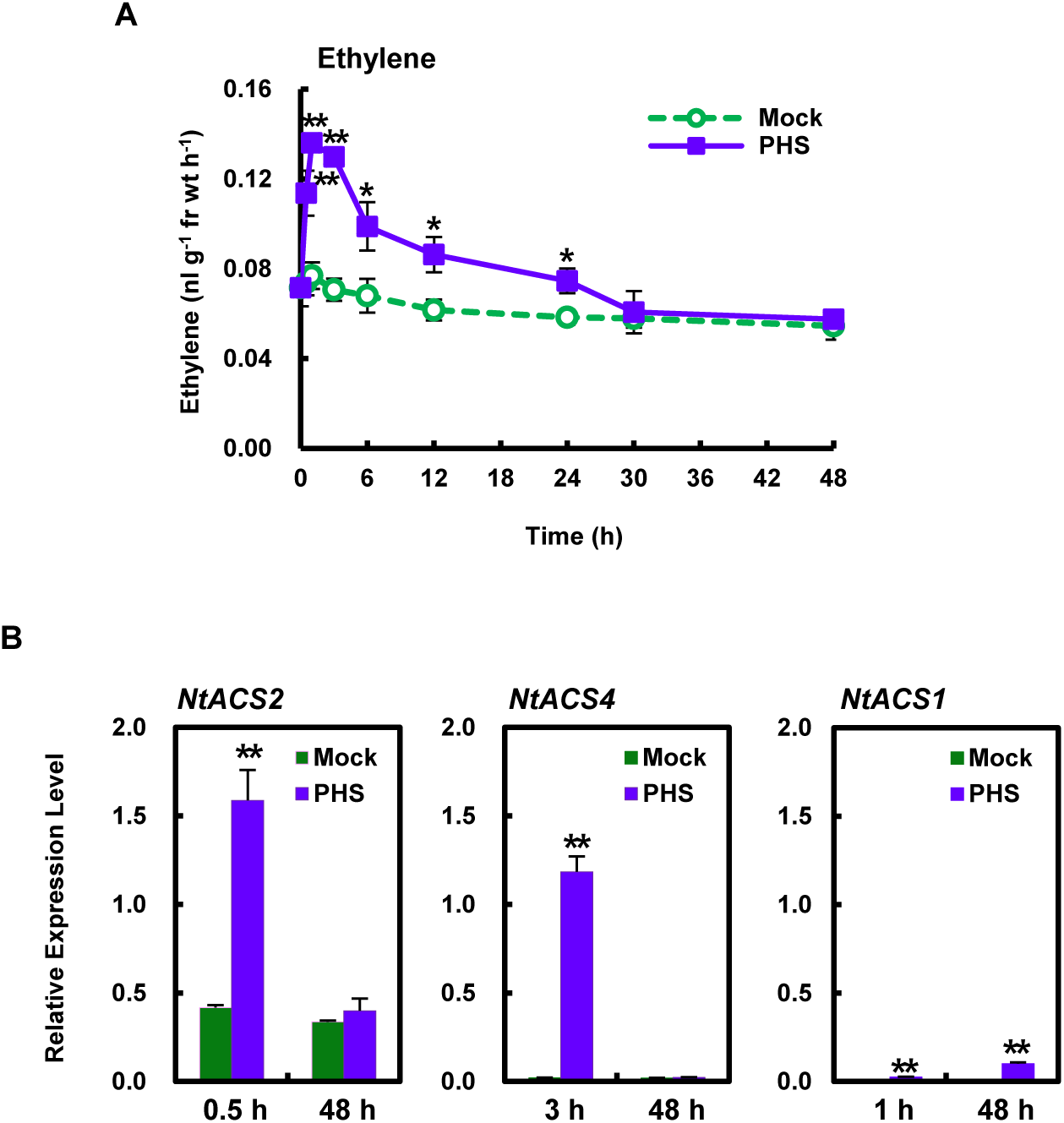
Kinetics of ethylene production and expression profiles of *ACS* isoforms in response to PHS treatment. A, Ethylene production in tobacco leaves after treatment with 1 μM PHS. Whole leaves of tobacco plants were treated with 1 μM PHS for 48 h. B, Transcription levels of *NtACS* gene family members *NtACS2* (left panel), *NtACS4* (middle panel), and *NtACS1* (right panel) at the indicated times after treatment with 1 μM PHS in tobacco leaves. Transcription levels are expressed relative to the reference gene *β-actin* after qRT-PCR. Ethylene levels and relative mRNA expression levels are expressed as means ± SD. An asterisk indicates a significant difference between mock- and PHS-treated cases (one asterisk (P < 0.05) or two asterisks (P < 0.01) at the same indicated time).

Abiotic stresses are known to induce ethylene production through gene-specific expression of 1-aminocyclopropane-1-carboxylic acid (ACC) synthase (ACS) members in a time-dependent manner (Wi et al., 2010). Apoptosis-related *NtACS2* and *NtACS4* expressions have been shown to induce early ethylene production, as early as after 1 h, in response to either abiotic/oxidative or biotic stress induced by the compatible pathogen *Ppn*, whereas necrosis-or senescence-related *NtACS1* expression was shown to be responsible for late-phase ethylene production at 24 h or 72 h for induction of oxidative or biotic stress, respectively (Wi et al., 2010; 2012).

In this study, we measured the transcription levels of *NtACS1, NtACS2*, and *NtACS4* in response to PHS treatment by performing qRT-PCR. Although *NtACS2* transcription was detected at a relatively low level in mock-treated plants at 0.5 h and 48 h, it was up-regulated by about 3.8-fold at 0.5 h in PHS-treated tobacco plants compared with that in mock-treated plants (Fig. 3B). However, it returned to the basal level at 48 h, at which time there was no significant difference in *NtACS2* expression between PHS- and mock-treated plants. On the other hand, *NtACS4* expression was remarkably up-regulated, by approximately 59-fold, at 3 h compared with mock-treated tobacco plants. However, apoptosis-related *NtACS4* transcription was almost absent at 48 h in both mock- and PHS-treated plants. These results are consistent with our previous finding that apoptotic-like cell death, which was dependent on *NtACS2* and *NtACS4*, was not further induced at 48 h after PHS treatment.

We previously reported that expression of *NtACS1*, a senescence- or necrosis-related *ACS* gene member, increased starting at 30 h and peaked at 72 h after compatible pathogen inoculation, indicating *NtACS1* is active in plant death during the later phase of pathogen infection and has a role different from those of *NtACS2* and *NtACS4* (Wi et al., 2012). PHS-induced *NtACS1* transcription was insignificantly increased at 1 h and 48 h (Fig. 3B). Therefore, based on the low levels of *NtACS1* transcription and ethylene production at 48 h after PHS treatment, PHS-induced cell death was not accompanied by necrosis-related plant death. This observation corresponds well with the monophasic production patterns of ROS and ethylene at an early stage following PHS treatment, indicating that additional necrotic cell death did not occur at a later stage (Fig. 1A; Fig. 2, A and B; Fig. 3A). These results further suggest that cell death induced by PHS is related to the early massive increases in ROS production and signaling for resistance induction.

### Effect of PHS on pathogen growth and cell damage in response to infection with a hemibiotrophic pathogen

In order to investigate why cell damage was not serious following PHS treatment, we examined the effects of PHS on pathogenicity after infection with the hemibiotrophic pathogen *Phytophthora parasitica*, which induces biotrophic activity during early infection and necrotrophic activity in the later stage of colonization (Shibata et al., 2010).

PHS-treated tobacco plants did not develop additional cell damage even until 120 h in culture bottles containing agar (Fig. 4A). Therefore, we first investigated the effect of pathogen-induced cell damage in the presence of PHS at the necrotic stage up to 120 h after pathogen infection. Hemibiotrophic pathogen *Ppn* gradually increased cell damage in tobacco plants, whereas PHS treatment significantly alleviated cell damage following pathogen invasion until 120 h, compared to that in plants treated only with pathogens (Fig. 4B). Trypan blue staining results revealed that PHS treatment did not further aggravate cell damage between 3 h and 120 h after pathogen infection, implying the presence of a resistance mechanism working against the pathogen infection.

**Figure 4.**
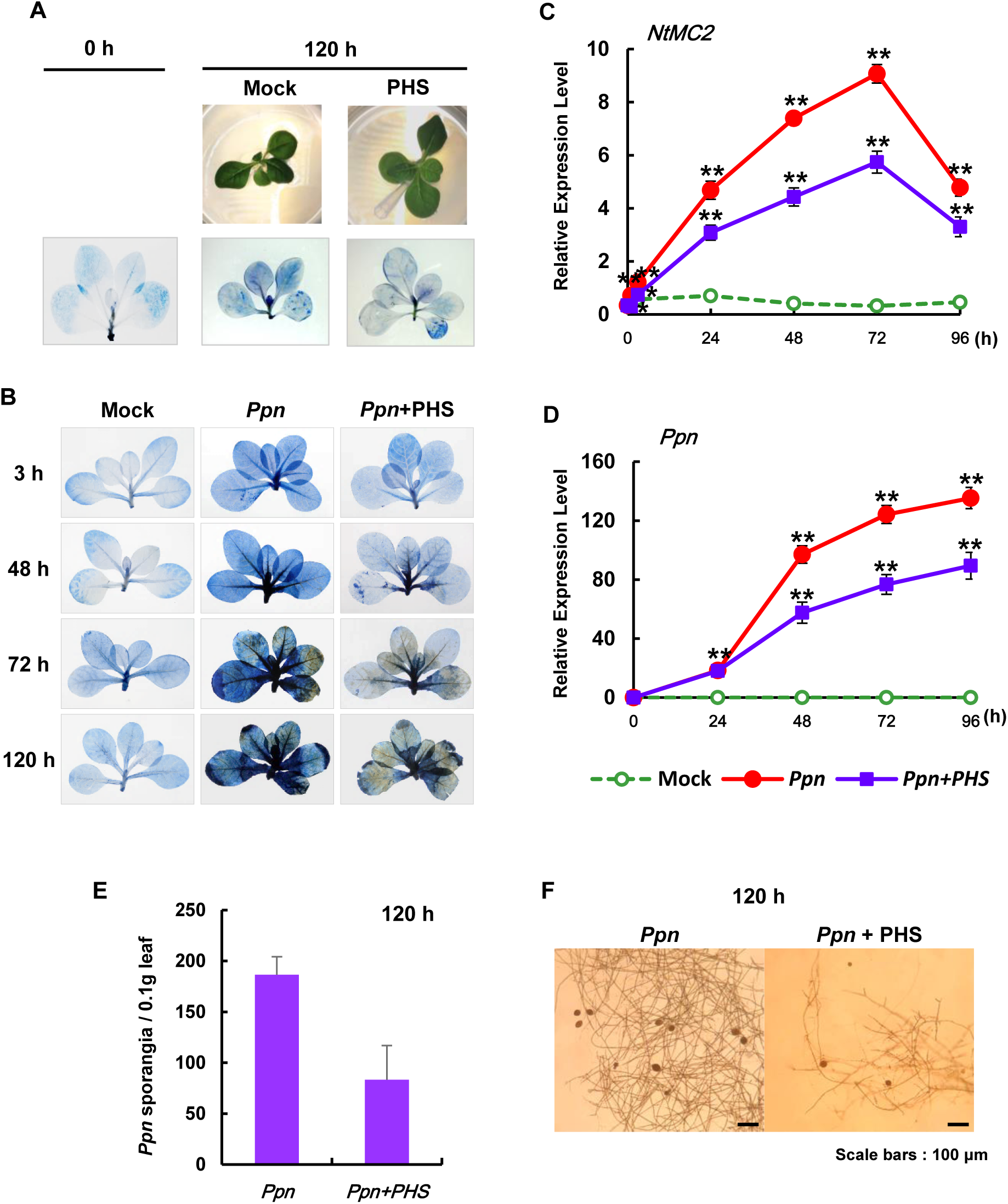
Effect of PHS treatment on cell damage and pathogen growth in *Ppn*-inoculated tobacco leaves. A, Necrotic areas in tobacco plants were determined after 1 μM PHS treatment using trypan blue staining and then imaged with a digital camera. Cell damage was detected after treatment with 100 μL of 1 μM PHS was slowly dropped on the shoot area using a micropipette tip. First line: mock treatment; Second line: photograph of plants treated with PHS for 120 h in culture bottles; Third line: cell damages in tobacco plants after treatment with PHS after trypan blue staining. B, Necrotic areas in *Ppn*-inoculated tobacco plants with or without 1 μM PHS treatment were determined using trypan blue staining and then imaged with a digital camera. Cell damage was detected at the indicated time after shoot inoculation with *Ppn* until 120 h, during which 100 μL of 1 μM PHS. C, Transcription levels of the tobacco *NtMC2* gene in *Ppn*-inoculated tobacco plants after PHS treatment for 96 h. Real-time qRT-PCR analysis of *NtMC2* transcription was performed using total RNAs from pathogen-infected tobacco leaves with PHS treatment. D, Detection of *Ppn* in wild-type plants after inoculation by real-time qRT-PCR of the 5.8S rRNA of *Ppn* using total RNAs from pathogen-infected tobacco leaves with PHS treatment. Transcription levels are expressed relative to the reference gene *β-actin* after qRT-PCR. Relative mRNA expression levels are expressed as means ± SD. Asterisks indicate a significant difference in PHS-treated or co-treated cases with PHS and *Ppn* infection from mock-treated cases at the same time point (one asterisk (P < 0.05) or two asterisks (P < 0.01)). E-F, Determination of the number of sporangia in *Ppn*-infected tobacco leaves with or without PHS treatment. (E) The number of sporangia was counted from 0.1 g of tobacco leaves after 120 h of *Ppn* inoculation. (F) The hypha and sporangia were photographed in *Ppn*-infected tobacco leaves with (right) or without (left) PHS treatment.

The transcription level of *metacaspase 2* in *N. tabacum* (*NtMC2*), which is a fundamental part of the cell death machinery in response to abiotic and biotic stresses (Watanabe and Lam, 2011), was significantly induced, reaching a peak at 72 h after *Ppn* inoculation. PHS treatment significantly inhibited *Ppn*-induced transcription of *NtMC2*, suggesting that PHS contributes to reducing the pro-cell death role of metacaspase (Fig. 4C).

Real-time qRT-PCR is a reliable technique for the detection and quantification of plant pathogens, and it is increasingly being used in plant pathology investigations. We recently reported that qRT-PCR is a highly sensitive method for the quantification of *Ppn* in plants (Wi et al., 2012). We designed primers specific to 5.8S rRNA (GenBank AY769953) as an internal control for monitoring *Ppn* growth. Using qRT-PCR, expression of the 5.8S rRNA gene was compared to that of tobacco *β-actin*, which was used as the internal plant reference gene. Pathogen growth was continuous until 96 h, revealing an S-shaped growth curve after *Ppn* inoculation, which was significantly inhibited by PHS treatment; moreover, the inhibitory effect was more prominent during the necrotrophic stage (Fig. 4D).

Next, we determined the numbers of sporangia in tobacco leaves after *Ppn* infection with or without PHS treatment. After 120 h of *Ppn* infection, PHS significantly reduced the numbers of *Ppn*-produced sporangia (Fig. 4E). In addition, PHS significantly inhibited the growth of mycelium with sporangia from *Ppn* inoculates in tobacco leaves (Fig. 4F). Taken together, the results are in agreement with the observed abrogation of cell damage and pathogen growth upon co-treatment with pathogen and PHS, suggesting the presence of PHS-related pathogen resistance.

### PHS-induced activation of SphK and alteration of sphingolipid metabolism under virulent pathogen infection

Until recently, only a few studies have investigated the physiological role of PHS1P in plants. Both S1P and PHS1P are abundant in plants, but PHS1P is present in lesser amounts in animals (Pata et al., 2010). Both S1P and PHS1P regulate stomatal apertures, via regulation of ABA signaling, and guard cell turgor in plants, suggesting that phosphorylated sphingolipid metabolites are stress-related signaling messengers in plants (Coursol et al., 2005).

Therefore, we focused on assessing the metabolism of sphingolipids, especially SphK, which phosphorylates PHS to PHS1P. SphK also phosphorylates sphingosine and other sphingoid LCBs. We first determined the transcription level of *SphK* after PHS treatment and under pathogen infection by performing qRT-PCR. The mRNA level of *SphK* was significantly elevated upon PHS treatment in *Ppn*-inoculated tobacco plants compared with that in mock-treated controls (Fig. 5A). After PHS treatment, transcription of the *SphK* gene continuously increased until 96 h, and PHS was more effective from 48 h to 96 h during the necrotic stage of hemibiotrophic pathogen infection.

**Figure 5.**
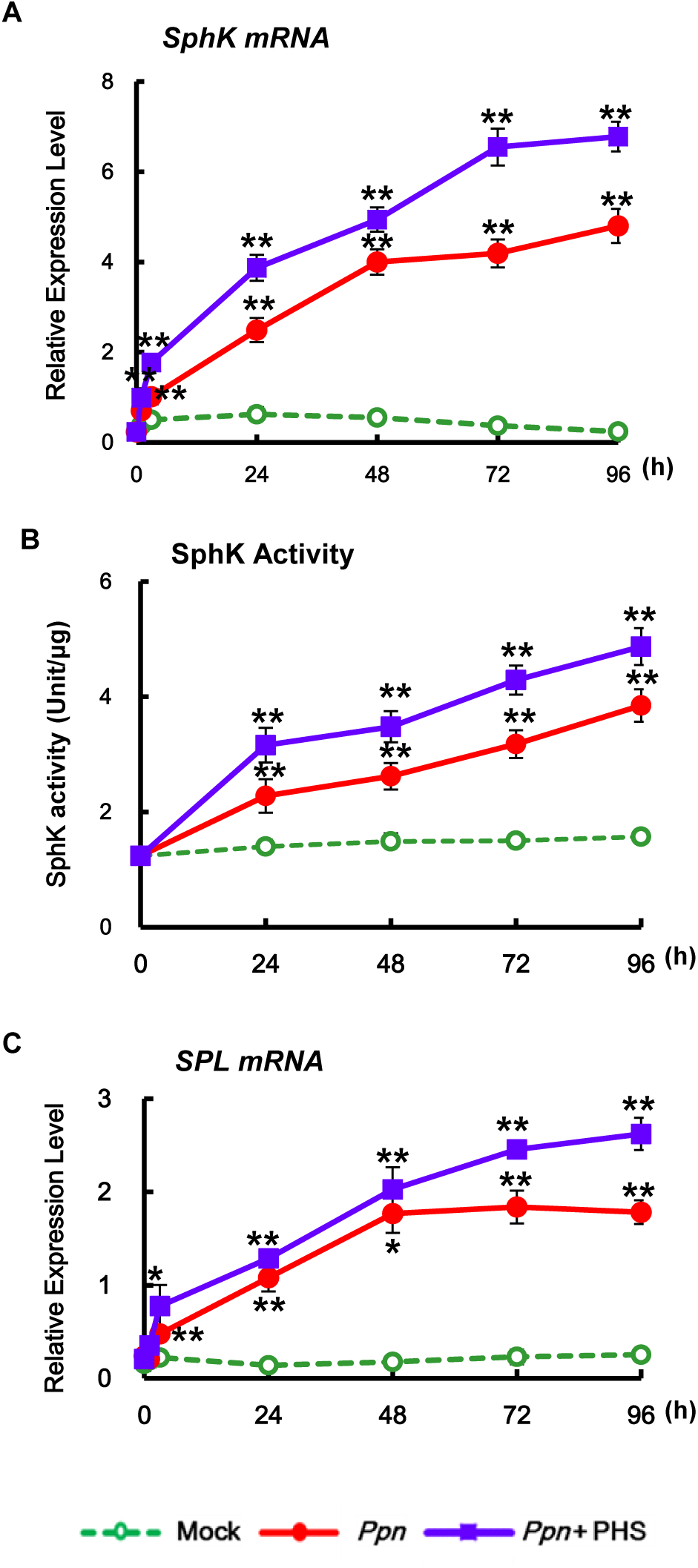
Effects of PHS treatment on the transcription and activity of sphingosine kinase. Effects of PHS treatment on the transcription and activity of sphingolipid metabolism. Relative transcription levels of SphK, SPT, and ceramidase in response to PHS treatment. Transcription levels of the *SphK* and *SPL* were determined in *Ppn*-inoculated tobacco leaves after treatment with 1 μM PHS for 96 h. A and C, Real-time qRT-PCR analysis of *SphK* (A) and *SPL* (C) transcription was performed in total RNAs from pathogen-infected tobacco leaves with PHS treatment. B, The activity of SphK was determined in *Ppn*-infected WT tobacco leaves with PHS treatment for 96 h. Asterisks indicate a significant difference in PHS-treated or co-treated cases with PHS and *Ppn* infection from mock-treated cases at the same indicated time (one asterisk (P < 0.05) or two asterisks (P < 0.01)).

PHS treatment further elevated the pathogen-induced increase of SphK activity in tobacco plants during the entire treatment period, compared with that in mock-treated leaves (Fig. 5B). The increases in the patterns of *SphK* gene expression and enzyme activity induced by PHS treatment were similar, suggesting that they are regulated at the transcriptional stage. The observation that up-regulation of *SphK* gene expression and SphK activity might be related to the attenuation of necrotic cell death by PHS at the later stage of *Ppn* treatment (48 h) suggests that PHS can act as a signaling molecule against pathogen attack through the activation of *SphK* transcription in plants.

In accordance with the SphK activity change, the enzyme responsible for degradation to PHS1P or S1P was more activated in PHS-treated tobacco plants after 48 h of *Ppn* infection, which is during the necrotic stage of the hemibiotrophic pathogen (Fig. 5C). These results are consistent with those in previous reports showing that an increase in S1P causes disease resistance in necrotrophic pathogens (Magnin-Ronert et al., 2015).

We next determined whether alteration of sphingolipid metabolism occurs after PHS treatment in *Ppn*-infected tobacco plants. LCB molecules, which are unique components of sphingolipids, are formed by SPT. As SPT catalyzes the first step of sphingolipid biosynthesis, gene expression levels of the LCB1 and LCB2 subunits were determined in order to elucidate sphingolipid metabolism after PHS treatment under *Ppn* infection (Fig. 6). Although transcription of *LCB1* and *LCB2* was unaltered in mock-treated control plants, transcription levels of both genes were significantly up-regulated from 24 h after *Ppn* inoculation, with higher expression of *LCB2* than *LCB1*. PHS treatment resulted in further significant elevations of *LCB1* and *LCB2* transcription during the entire period of pathogen infection; PHS-induced activation had the strongest effect on LCB1 at 72 h and on LCB2 at 48 h (Fig. 6, A and B). However, *ORM1* and *ORM2* transcription rapidly increased after PHS treatment (Fig. 6, C and D), suggesting the rapid increase is responsible to sphingolipid homeostasis. These results indicate that the *de novo* synthesis of sphingolipids occurs through feedback activation that is induced by exogenously added PHS in *Ppn*-infected tobacco plants. PHS-induced co-activations of *LCB1, LCB2*, and *SphK* transcription is responsible for metabolizing to PHS1P or S1P, which are bioactive sphingolipids that regulate many cellular responses (Proia and Hla, 2015). These observations indicate that sphingolipids are induced in response to pathogen attack, and PHS is an effective molecule in pathogen-induced sphingolipid biosynthesis.

**Figure 6.**
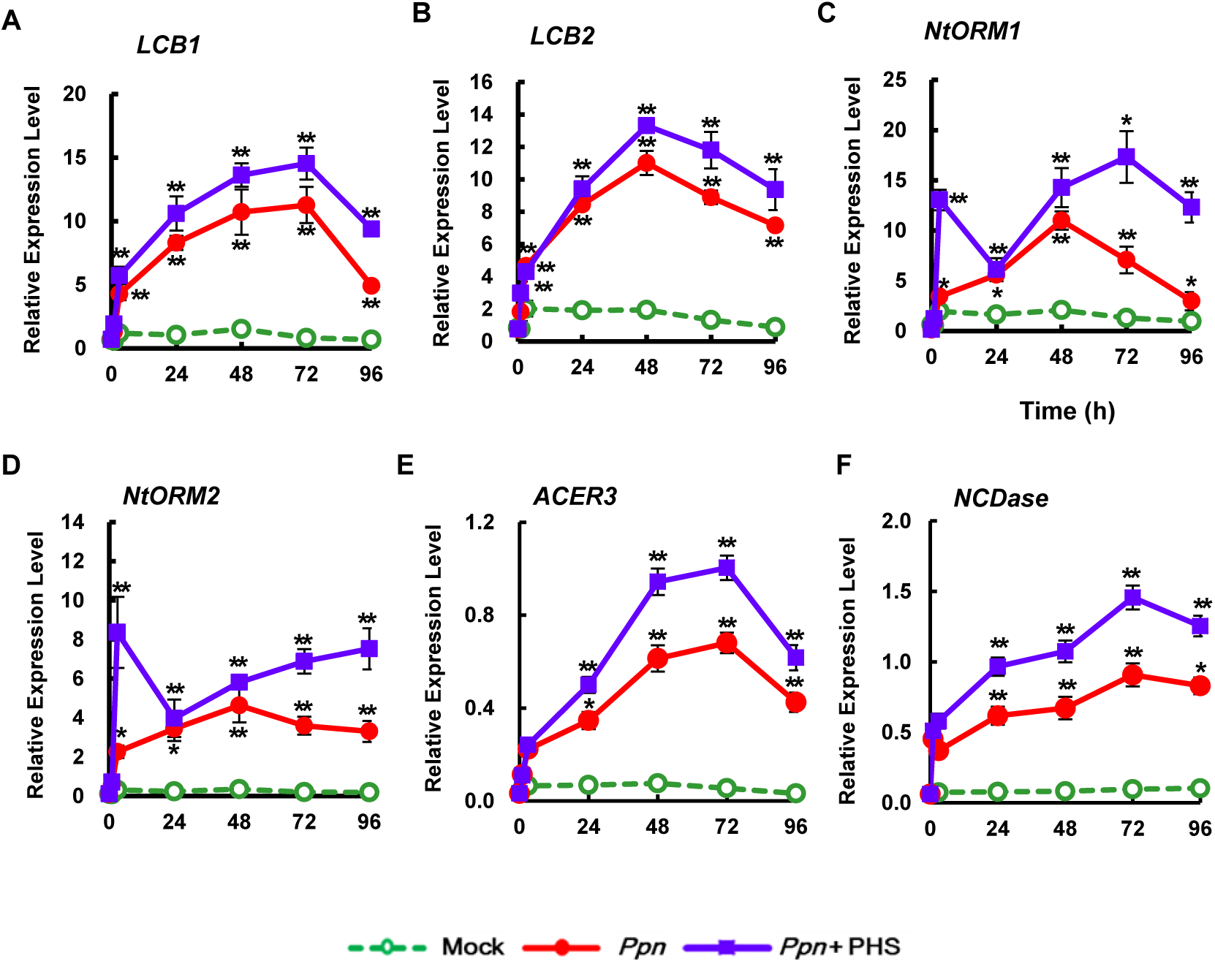
Effects of PHS treatment on sphingolipid metabolism. A-B, Transcription levels of two subunits, *LCB1* (A) and *LCB2* (B), of SPT were determined for 96 h after PHS treatment. C-D, Transcription of two genes, *ORM1*(C) and *ORM2* (D), of orosomucoid were determined after PHS treatment. Real-time qRT-PCR analysis of each gene was performed using total RNAs from pathogen-infected tobacco leaves with PHS treatment. E-F, Transcription levels of ceramidases *ACER3* (E) and *NCDase* (F) were determined for 96 h after PHS treatment. Transcription levels are expressed relative to the reference gene *β-actin* after qRT-PCR. Relative mRNA expression levels are expressed as means ± SD. Asterisks indicate a significant difference in PHS-treated or co-treated cases with PHS and *Ppn* infection from mock-treated cases at the same time point (one asterisk (P < 0.05) or two asterisks (P < 0.01)).

PHS influences the composition of sphingolipid metabolites in the next steps of SPT was further investigated. Transcription of alkaline ceramidase (*ACER3*), which hydrates unsaturated ceramides, dihydroceramides, and phytoceramides to form LCBs such as PHS or sphingosine (Mashima et al., 2020), was effectively elevated, by approximately 63%, after 48 h of PHS treatment compared to that after *Ppn* inoculation (Fig. 6E). Moreover, transcription of neutral ceramidase (*NCDase*), which may protect against apoptotic cell death by preventing the accumulation of ceramides in cells (Osawa et al, 2005), was elevated by about 53% at 48 h after co-treatment with *Ppn* and PHS (Fig. 6F). Collectively, these results suggest that PHS-induced elevation of *LCB1, LCB2, ACER3*, and *NCDase* transcription in accordance with increased *SphK* expression and SphK activity can contribute to attenuation of pathogen-induced necrotic cell death at a later stage.

### PHS-induced expression profiles of ROS-detoxifying enzymes and *PR* genes under pathogen infection

Next, the effects of PHS on ROS generation were determined in tobacco plants during the necrotic stage of hemibiotrophic pathogen infection. We previously reported a second massive ROS peak indicating necrosis between 36 h and 48 h in a compatible interaction with *Ppn*, in which pathogen growth is extensive (Wi et al., 2012). During the necrotic stage under a compatible *Ppn* pathogen infection, PHS treatment inhibited ROS accumulation by approximately 40% at 48 h and reduced the transcription of *NtRbohD* and *NtRbohF* in *Ppn*-inoculated tobacco plants (Fig. 7; Supplemental Fig. S2). This result is contrary to the PHS-induced up-regulation of *NtRbohD* transcription in the HR-relevant apoptotic stage at 1 h without *Ppn* infection (Fig. 2C). Although PHS had an inhibitory effect on the second ROS peak, it had little effect on the first ROS peak under *Ppn* infection, indicating PHS exhibits its function in a time-dependent manner. In addition, PHS did not affect *Ppn*-induced expression of *NtACS2* and *NtACS4* for ethylene production at the early stage, but slightly reduce that of *NtACS1* at the later stage (Supplemental Fig. S3). ROS production at a later stage might be a byproduct or harmful substance of cell damage in the necrotic stage of compatible pathogen infection.

**Figure 7.**
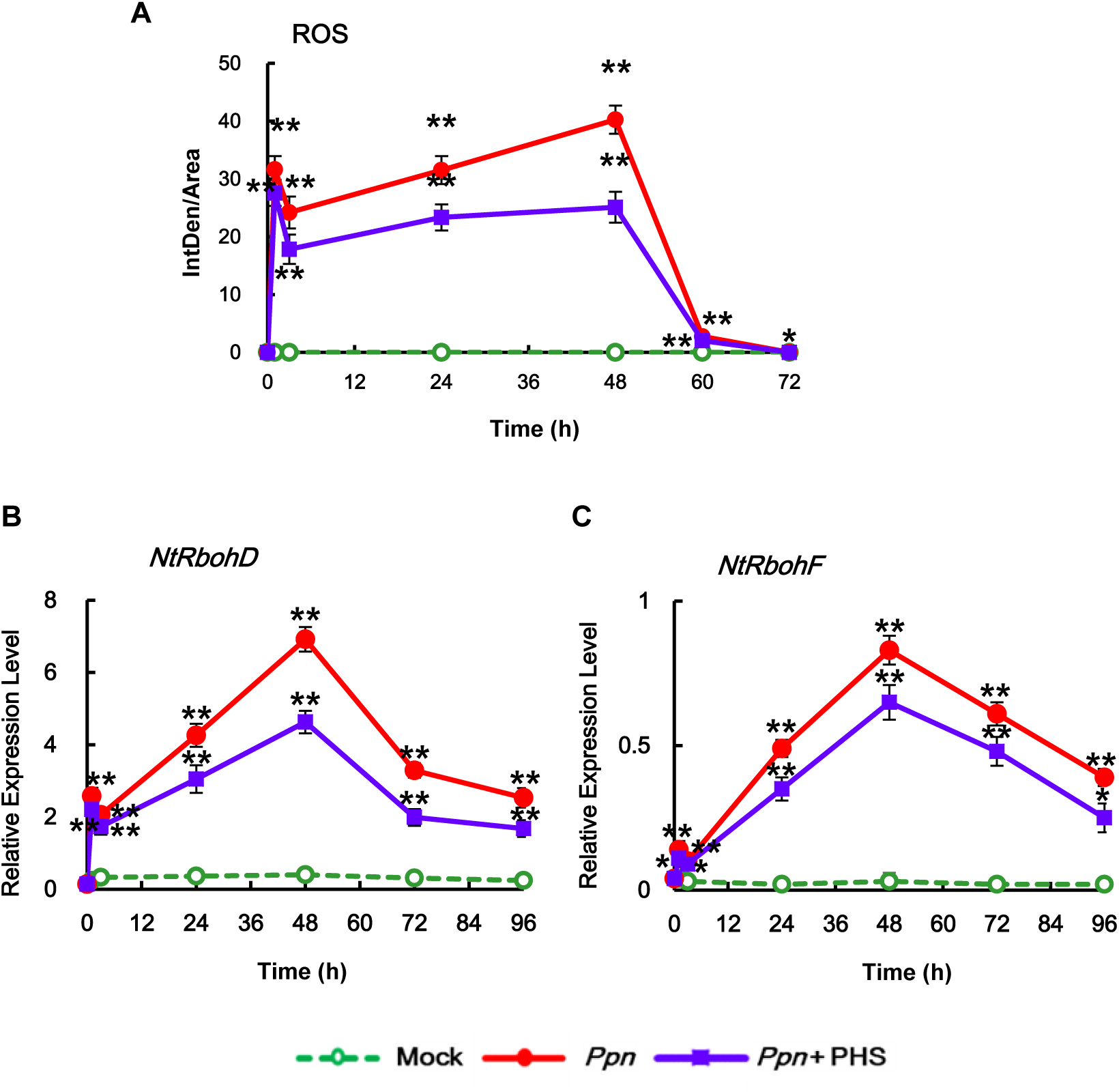
Effects of PHS treatment on ROS accumulation and *NADPH oxidase* gene (*NtRbohD* and *NtRbohF*) transcription in pathogen-infected tobacco leaves. A, After histochemical analysis of intracellular ROS accumulation in *Ppn*-infected tobacco leaves after PHS treatment, the intensity of fluorescence was quantified by ImageJ software. After mature tobacco leaves were treated with 1 μM PHS for the indicated time, ROS was determined by incubation with DCFH-DA for 10 min. Staining images of leaves were obtained by confocal microscopy and then quantified by ImageJ software. B-C, Relative mRNA levels of *NtRbohD* and *NtRbohF* genes in mature tobacco leaves infected with *Ppn* and then treated with PHS. Transcription levels of *NtRbohD* (B) or *NtRbohF* (C) are expressed as means ± SD. Transcription levels are expressed relative to the reference gene *β-actin* after qRT-PCR. Relative mRNA expression levels are expressed as means ± SD. Asterisks indicate a significant difference in PHS-treated or co-treated cases with PHS and *Ppn* infection from mock-treated cases at the same indicated time (one asterisk (P < 0.05) or two asterisks (P < 0.01)).

Therefore, we investigated whether PHS contributes to the expression of ROS-detoxifying enzymes during the later stage of *Ppn* infection. An enzymatic dismutation reaction converts superoxide into a more stable, membrane-permeable hydrogen peroxide (H_2_O_2_) derivative, which is required for cell-to-cell signaling. ROS-scavenging enzymes such as superoxide dismutase (SOD), ascorbate peroxide (APX), and catalase (CAT) provide cells with highly efficient mechanisms for detoxifying superoxide and H_2_O_2_ (Foyer and Noctor, 2005).

PHS treatment markedly up-regulated the gene expression of enzymes involved in ROS-detoxifying pathways, including mitochondrial manganese-SOD (MnSODmi), cytosolic copper/zinc SOD (CuZnSODc), cytosolic APX (APXc), CAT (CAT1 and CAT2), and phi glutathione-S-transferase (GSTF) (Fig. 8). *CAT1* and *CAT2* expressions were induced biphasically at 24 h and 72 h after *Ppn* infection and were significantly enhanced by PHS treatment (Fig. 8, D and E). On the other hand, expressions of other ROS-detoxifying enzymes were significantly induced monophasically in the later stage of PHS treatment under *Ppn* infection (Fig. 8, A, B, and C). PHS treatment up-regulated transcription of the SODs *MnSOMmi* and *CuZnSODc*, reaching maximum peaks at 72 h and 48 h, respectively, and resulted in an elevation of *APXc* transcription, which peaked at 48 h for detoxification of H_2_O_2_, compared with that in only *Ppn* infection. Expression of *GSTF*, a plant-specific phi class of stress-induced GSTs, reached a peaked at 72 h under *Ppn* inoculation, which was elevated further by PHS treatment (Fig. 8F). Therefore, it can be suggested that PHS activates the ROS detoxification pathway, which functions in lowering ROS accumulation, resulting in the prevention of severe necrosis at a later stage of compatible pathogen infection. Therefore, it is suggested that PHS participates in plant survival, or rather, it prohibits further progress of cell damage in a time-dependent manner suggests.

**Figure 8.**
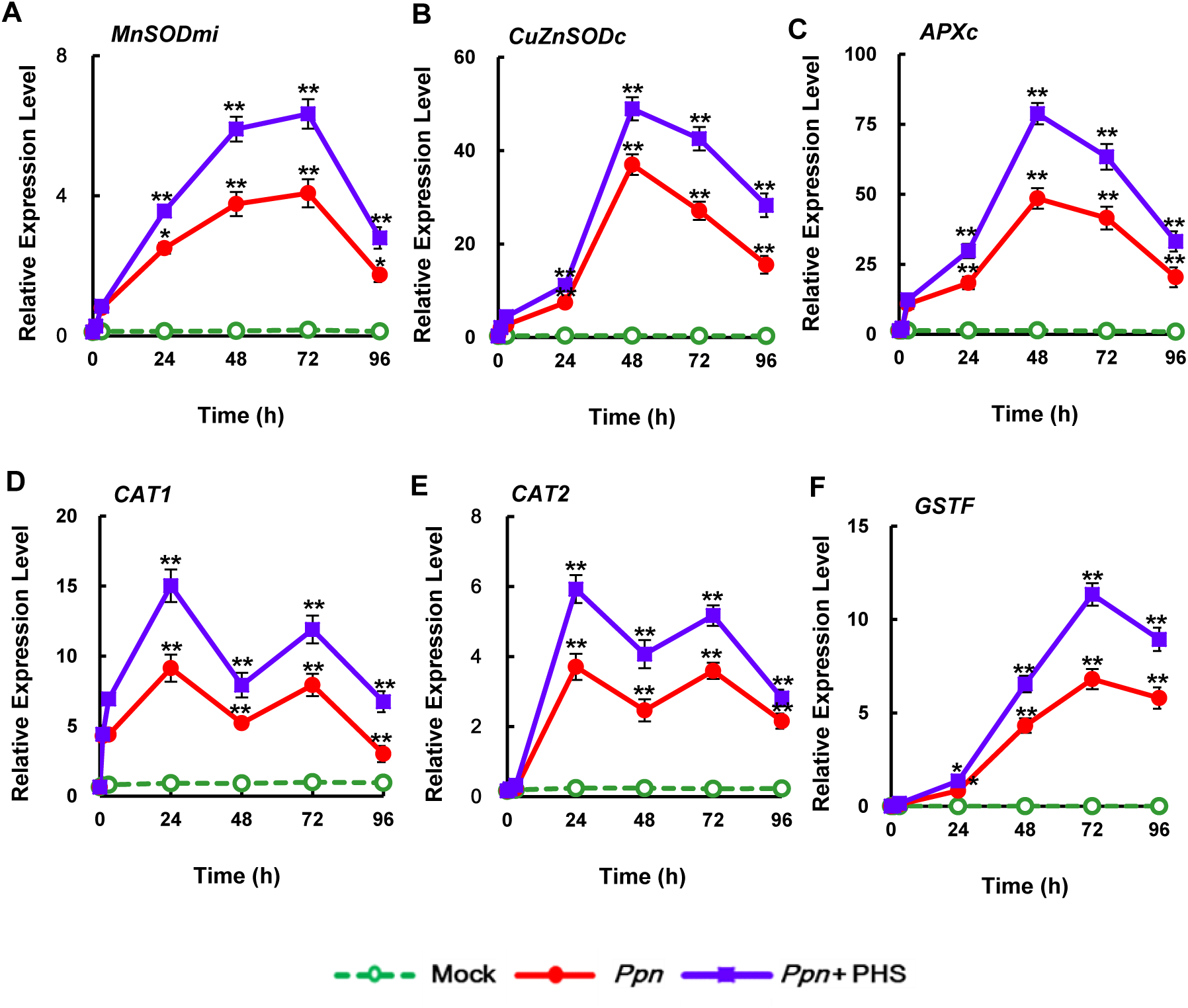
Relative transcription levels of endogenous ROS detoxification enzymes, *CAT1, CAT2, MnSODmi, CuZnSODc, APXc*, and *GSTF*, after 1 μM PHS treatment for 96 h in leaves of tobacco plants. PHS-treated whole leaves were subjected to real-time qRT-PCR analysis. Transcription levels are expressed relative to the reference gene *β-actin* after qRT-PCR. Relative mRNA expression levels are expressed as means ± SD. Asterisks indicate a significant difference in PHS-treated or co-treated cases with PHS and *Ppn* infection from mock-treated cases at the same time point (one asterisk (P < 0.05) or two asterisks (P < 0.01)).

We next investigated whether PHS can affect the expression of *PR* genes, which are responsible for the defense response against pathogen infection. As shown in Fig. 9, PHS treatment significantly elevated mRNA levels of *PR-1a, PR-3, PR-4b, PR 5* (*OSM*), and *SAR 8.2* at 48 h, as well as *PR 5* and *nonexpressor of PR1* (*NPR1*) at 72 h in pathogen-infected tobacco plants compared with that in pathogen infection alone plants. In addition, transcription patterns of *PR* genes were characterized by a transient elevation followed by a gradual reduction to significantly lower levels after 96 h under the compatible pathogen infection. Transcriptions of *osmotin-like* (*OSM*) and *taumatin-like* (*TLP*) proteins, which belong to the *PR5* family, were more markedly up-regulated by PHS treatment after 48 h and 72 h of *Ppn* inoculation, respectively, suggesting enhancement of the PHS effect on pathogen tolerance (Fig. 8). Transcription of *SAR8.2*, which has antifungal activity (Lee and Hwang, 2006), was most significantly elevated at 48 h after PHS treatment under pathogen infection, indicating that PHS possesses effective antifungal activity in response to fungal pathogens. These results indicate that up-regulation of *PR* gene expression is responsible for PHS-induced attenuation at the later stage of pathogenesis.

**Figure 9.**
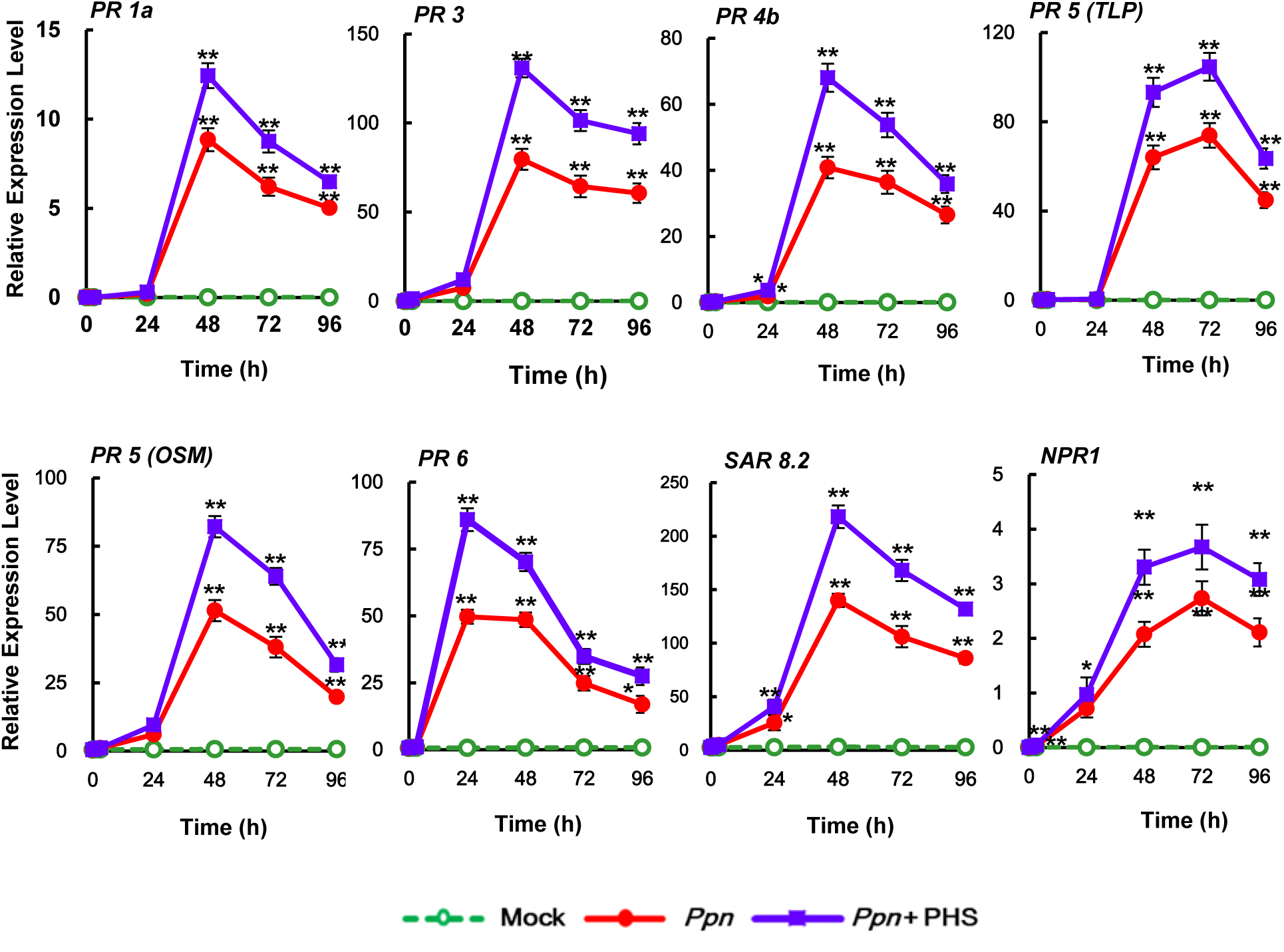
Effects of PHS treatment on the transcription of *PR* genes. Relative transcription levels of PR proteins under pathogen infection in response to PHS treatment for 96 h. Transcription levels of the *PR* genes *PR-1a, PR-3, PR-4b*, taumatin-like proteins (*TLP*), and osmotin (*OSM*) for *PR5, PR-6, SAR8.2*, and *NPR1* were determined in tobacco leaves after treatment with 1 μM PHS. Transcription levels are expressed relative to the reference gene *β-actin* after qRT-PCR. Relative mRNA expression levels are expressed as means ± SD. Asterisks indicate a significant difference in PHS-treated or co-treated cases with PHS and *Ppn* infection from mock-treated cases at the same time point (one asterisk (P < 0.05) or two asterisks (P < 0.01)).

### Involvement of SphK in response to *Ppn* infection

We further investigated *SphK* gene expression and sphingolipid metabolism in comparative experiments between a virulent pathogen and the avirulent elicitor cryptogein. *SphK* transcription was rapidly induced and reached a maximum (3.2-fold) at 24 h after cryptogein treatment compared with that in pathogen-treated tobacco plants without cryptogein (Supplemental Fig. S4). Following this, the level of *SphK* transcription gradually decreased. In contrast, *SphK* transcription remained low in susceptible tobacco plants after *Ppn* infection until 24 h and gradually increased thereafter. We further investigated whether ROS are a determinant of *SphK* transcription during the biotic stress response. Using *RbohD-AS* and *RbohF-AS* transgenic tobacco plants, in which ROS production is impaired by the antisense expression of *NtRbohD* and *NtRbohF, SphK* transcription was markedly enhanced only at 48 h following *Ppn* infection compared to that in wild-type (WT) plants. Transcription levels of *SphK* in *Ppn*-infected *RbohD-AS* and *RbohF-AS* plants increased by 5.6- and 2.7-fold, respectively, compared to that in *Ppn*-infected WT plants after 48 h (Supplemental Fig. S5). Higher activation of *SphK* transcription in *RbohD-AS* and *RbohF-AS* plants suggests a greater role for lower pathogenicity in response to virulent pathogen infection.

To further determine whether ethylene affects *SphK* expression, *SphK* transcription was measured in *CAS-AS* transgenic plants, in which ethylene production is impaired by the antisense expression of the ACC synthase gene (Wi et al., 2012). However, *SphK* transcription levels in *Ppn*-infected *CAS-AS* plants were similar to those in pathogen-infected WT tobacco plants, indicating that ethylene is not a determinant of *SphK* expression.

Next, LCBPs treatment was performed to investigate the effects on *Ppn*-induced cell damage. LCBPs were treated with 1 μM of S1P or PHS1P along with *Ppn* pathogens on tobacco plant stems with about 5 leaves. Both S1P and PHS1P treatment inhibited cell damage effectively in plants at 24 h and 72 h after pathogen infection (Fig. 10). Despite the administration of PHS1P and S1P at the base of the stem, cell damage to the leaves at the top of the stem was remarkably weak. The results indicated that PHS1P and S1P can induce SAR, increasing the resistance of the whole plant. Taken together, these observations indicate the possibility that SphK might be involved in preventing necrosis at the beginning of the necrotic stage.

**Figure 10.**
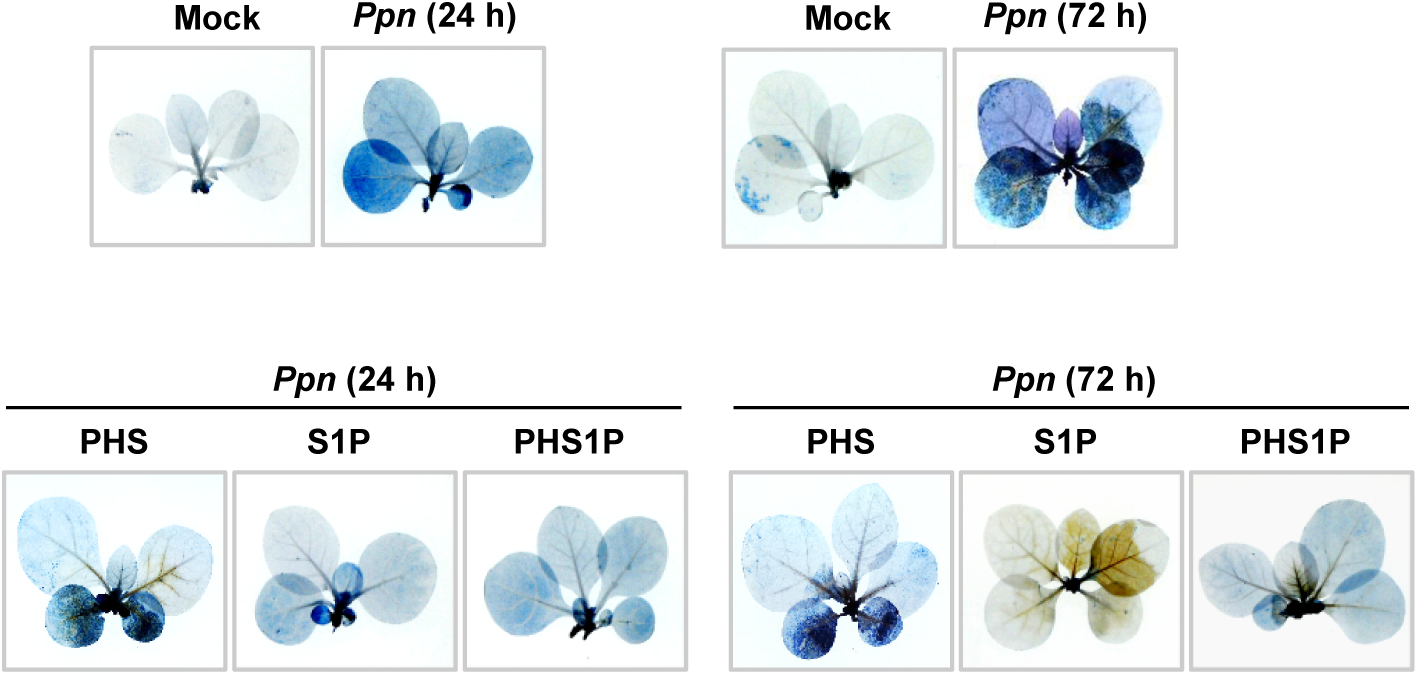
Effect of S1P and PHS1P treatment on cell damage in *Ppn*-inoculated tobacco leaves. Necrotic areas in *Ppn*-inoculated tobacco plants were determined after 1 μM of S1P or PHS1P treatment by using trypan blue staining and imaging with a digital camera. Cell damage was detected after shoot inoculation with *Ppn* for 24 h and 72 h, during which 100 μL of 1 μM PHS was slowly dropped on the shoot area using a micropipette tip.

## Discussion

Sphingolipids, a class of well-known lipids in mammal cells, have recently been shown to have important functions in plants (Aguilera-Romero et al., 2013). The mono-unsaturated dihydroxy LCB (d18:1) and the saturated or mono-unsaturated trihydroxy LCB (t18:0 and t18:1, respectively) are the main LCBs in plants (Cacas et al., 2012). Although the *cis* (*E*) and *trans* (*Z*) isomers of 4-hydroxy-8-sphingenine (t18:1) comprise the major forms in *Arabidopsis* and tomato (Dietrich et al., 2008), PHS (t18:0) is a major LCB in tobacco plants, and there are quite similar LCB compositions in *N. benthamiana* and *N. tabacum* leaves (Cacas et al., 2012).

Sphingolipids, as bioactive molecules, have been shown to have physiological functions in the stress responses and PCD in plants (Berkey et al., 2012). Sphingolipid accumulation is an intrinsic early step in HR activation during the pathogen response. Sphingolipids derived from the fungal pathogen *Magnaporthe grisea* have been shown to induce phytoalexin accumulation, PCD, and resistance to infection in rice (Koga et al., 1998). It was recently reported that sphingolipids such as PHS and PHS1P have important roles in signal transduction pathways that regulate physiological functions and stress responses in plants, especially the immune response (Rivas-San Vicente et al., 2013).

In *N. benthamiana*, sphingolipid biosynthesis is crucial during the early stages of defense responses to the non-host pathogen *Pseudomonas cichorii* as well as the host pathogen *P. syringae* pv. *tabaci* (Takahashi et al., 2009). We previously reported an elevation of PHS in response to the virulent pathogen *Ppn* in tobacco leaves after 1 h and 48 h (Cho et al., 2012). It is suggested that PHS is a beneficial regulator during pathogenesis (Berkey et al., 2012). Therefore, it can be suggested that the rapid accumulation of sphingolipid bases such as sphingosine and PHS is an important step for the resistance response against pathogen attack. However, exactly how pathogens trigger PHS accumulation has not been described, and whether or not PHS accumulation has a role in the restriction of bacterial growth during the pathogen response also remains to be elucidated. Further, the physiological significance of PHS accumulation in pathogen-infected plants requires explanation. On that basis, our study focused on the pathophysiological significance of PHS.

Transcription and activation of SphK were markedly up-regulated at 96 h upon PHS treatment (Fig. 1, G and H). Therefore, it could be suggested that these characteristics were related to the abrogation of the second phase of ROS accumulation and necrotic cell death. As PHS and its phosphorylated form have been reported to mediate pathogen tolerance in several plants against necrotrophic pathogens (Magnin-Robert et al., 2015), these results suggest that PHS might be involved in the development of protective machinery against pathogen-induced cell death through conversion of PHS into other protective compounds such as PHS1P.

Recent evidence has indicated that phosphorylated sphingosines such as S1P, which is formed from sphingosine by SphK, can act as potent bioactive lipids for the survival of cancer cells (Riccitelli et al., 2013). Other recent studies have shown that SphK inhibition results in apoptosis in xenografts (Kapitonov et al., 2009) in animals. Further, exogenous S1P supplementation can propel murine splenocytes toward a pro-survival outcome in response to hypoxia-induced injury (Chawla et al., 2014). Moreover, PHS1P is known to increase the cell viability of human dermal fibroblasts via the c-Jun N-terminal kinase/Akt pathway (Lee et al., 2012).

It has also been reported that SphK is activated by ABA in *A. thaliana* for the inhibition of stomatal opening and the promotion of stomatal closure, suggesting that S1P is a signaling molecule involved in ABA regulation of guard cell turgor (Coursol et al., 2005). Recently, overexpression of the rice S1P lyase gene *OsSPL1* in transgenic tobacco was shown to result in reduced tolerance to salt and oxidative stresses compared to that in WT plants (Zhang et al., 2012). ROS generation induced by LCBs is specifically blocked by their phosphorylated forms, indicating that maintenance of homeostasis between a free sphingolipid base and a phosphorylated derivative is critical to determining cell fate (Shi et al., 2007). These observations, including our results (Fig. 10), likely indicate that enhancement of endogenous S1P and PHS1P positively regulates stress tolerance in plants.

In our results, the observation that PHS induced acute transient NADPH oxidase-dependent ROS accumulation and ethylene production only at an early stage are in contrast to the observed elevation of transcription and activation of SphK in a biphasic pattern under PHS treatment (Fig. 1, 2, and 3). PHS-induced rapid transient accumulation of ROS could prevent further progression of cell death, similar to that in the HR (Fig. 1B). This is supported by the observation that ROS at an early stage act as signaling molecules, signaling that can be triggered by PHS treatment.

In our study, expression of SphK was further activated by PHS in tobacco plants at a relatively later stage until 96 h compared to that in a *Ppn* only infection (Fig. 5, A and B). In the later stage of the *Ppn* infection, pathogen growth and pathogen-induced cell damage were significantly suppressed by PHS treatment (Fig. 4B-4F). These observations are consistent with the results showing that PHS1P and S1P can significantly suppress pathogen-induced plant damages (Fig. 10). Interestingly, alteration of the balance between PHS and phosphorylated PHS is commonly known to affect sphingolipid-mediated PCD or cell survival in plants and animals (Sánchez-Rangel et al., 2015). These results indicate that the magnitude of *SphK* transcription is related to the progression of plant tolerance against pathogenicity. Moreover, PHS up-regulated the expression of ROS-detoxifying enzymes, which may be responsible for decreasing ROS levels, and the expression of *PR* genes at the later stage of *Ppn* infection (Fig. 8 and 9). Elevation of *PR* gene expression resulted in disease resistance such as SAR in the necrotic stages of *Ppn* infection in PHS-treated tobacco leaves.

We previously reported that the PHS level was shown to rapidly increase (by 1.8-fold at 1 h and 2.36-fold at 48 h) following virulent pathogen inoculation in tobacco plants (Cho et al., 2012). In another study, the ceramide kinase mRNA level was up-regulated by about 5-fold at 24 h in plants infected with virulent *P. syrangae* compared to the level in an uninfected control (Liang et al., 2003). Considering our previous detection of a biphasic increase in PHS level (Cho et al., 2012) and the continuously increasing SphK activity up to 96 h (Fig. 5B) after *Ppn* infection, it is suggested that PHS is metabolized into phosphorylated PHS at the later stage under *Ppn* infection. Collectively, these observations indicate that PHS can act as a signaling molecule by activating *SphK* transcription for the suppression of necrotic cell death when infected by the hemibiotrophic pathogen *Phytophtora parasitica*. Therefore, based on the profile of *SphK* transcription, increased induction of SphK is notably advantageous to plant immunity.

*SphK* transcription was significantly increased in *RbohD-AS* and *RbohF-AS* transgenic plants (Supplemental Fig. S5), which have a much more tolerant phenotype against *Ppn* infection, indicating a negative correlation between the level of pathogenicity and *SphK* transcription at later stage (48 h), although ROS act as signaling molecules for a PHS-induced defense response at the early stage (Fig. 7A). Moreover, not only the magnitude but also the timing of ROS-independent *SphK* transcription were found to be important determinants of necrotic cell death in virulent pathogen-infected plants.

To the best of our knowledge, there have been few studies on the effects of ROS on sphingolipids or their metabolism in plants and animals. ROS production was detected relatively early after treatment with an S1P, implying that S1P acts as a signaling molecule through immediate ROS generation (Keller et al., 2006). However, inhibition or knockdown of SphK has been shown to enhance ROS formation and apoptosis in animal cells rat cardiac cells (Pchejetski et al., 2007). Further, monoamine oxidase-dependent ROS generation leads to mitochondria-mediated apoptosis along with SphK inhibition in rat cardiac cells (Pchejetski et al., 2007). The evidence confirming up-regulation of the ceramide/S1P ratio as a critical event in animal cell apoptosis suggests a link between SphK inhibition and ROS generation (Pchejetski et al., 2007).

In our system, ROS were generated in a biphasic manner under *Ppn* infection with the first peak at 1 h and the second peak at 48 h (Fig. 7A). In the second ROS peak, the period and amount of ROS generation were very high. It is thought that the second ROS peak causes cell damage and death unlike the first ROS peak, which acts as a signaling event. The amount produced at the second ROS peak was notably lower when the pathogen and PHS were treated together than when the pathogen alone was treated. These results indicate that PHS can alleviate pathogen-induced ROS accumulation and damage at the later stage. In particular, when PHS1P and S1P were treated to the tobacco stem, cell damages were significantly suppressed even in the leaves above the treatment point (Fig. 10), suggesting that these LCBPs can induce SAR in response to a pathogen attack.

In conclusion, these observations in this study suggest that elevated gene expressions of long-chain sphingolipid bases for *de novo* synthesis of a sphingolipid is an important determinant for preventing necrotic cell death in response to pathogen attack. These observations also suggest that an elevated ratio of phosphorylated/unphosphorylated long-chain sphingolipid bases induced by SphK may be a more significant determinant for promoting resistance in a time-dependent manner during plant-pathogen interactions. Therefore, the rapid induction of PHS is beneficial for the activation of SphK and inhibition of pathogenicity in the necrotic stage of a hemibiotrophic infection, resulting in the development of SAR in plant immunity. Taken together, our results indicate that the selective channeling of sphingolipids into their phosphorylated forms in conjunction with detoxification of stress-induced ROS and activation of *SphK* transcription has physiological pro-survival effects related to resistance in plants exposed to biotic stresses.

## Materials and Methods

### Plant Materials, Growth Conditions, PHS Treatment, and Fungal Inoculation

Surface-sterilized seeds of tobacco (*Nicotiana tabacum* L. Wisconsin 38) plants were cultured on solid Murashige and Skoog (MS) medium (pH 5.8) under a light cycle (16 h light/8 h dark, 100 μmol photons m^-2^ s^-1^) at room temperature (25 ± 5°C). Phytosphingosine (PHS) was obtained from Santa Cruz Biotech (Dallas, Texas, USA). Phytosphingosine-1-phosphate (PHS1P) and NBD sphingosine were purchased from Avanti Polar Lipids (Alabaster, Alabama, USA) and sphingosine-1-P (S1P) was purchased from Sigma-Aldrich (St. Louis, Missouri, USA. Solutions of PHS, PHS1P, and S1P with 20 mM MES buffer (pH 6.1) were applied to whole leaves through petioles or stems with five leaves. Tobacco shoots with four to five leaves were inoculated directly with a pathogen plug (1 cm diameter) in a culture bottle containing solid half-strength MS medium.

### RNA Isolation and Real-Time qRT-PCR

Total RNA isolation was performed as described previously (Seo et al., 2020). After 1 μg of total RNA from leaves was reverse-transcribed using a High Fidelity PrimeScript™ RT-PCR Kit (Takara, Japan), and Real-time qRT-PCR was performed using a Thermal Cycler Dice® Real Time System III T950 (Takara, Japan) with gene-specific PCR primers (Supplemental Table S1). Relative expression levels in each cDNA sample were normalized to the reference gene *β-actin*.

### Sphingosine Kinase Activity Assay

The activity of sphingosine kinase was measured by performing the fluorescence-based *in vivo* assay with NBD sphingosine (Pitman et al., 2017). To extract crude proteins from 0.3 g of tobacco leaves, frozen leaves were ground into powder and suspended in 300 μL of lysis buffer (50 mM Tris-HCl, pH 7.4, 150 mM NaCl, 10 % glycerol (w/v), 1 mM dithiothreitol, 2 mM Na_3_VO_4_, 10 mM NaF, 10 mM β-glycerophosphate, 1 mM EDTA and 0.5 mM phenylmethylsulfonyl fluoride). Then, 200 μL of protein extracts were incubated for 1 h at 37°C with 10 μM NBD sphingosine. The reaction was initiated by adding 10 μM ATP in final concentration to the reaction buffer. At the end of the reaction, the reaction was stopped by adding 270 μL of chloroform/methanol/HCl (100:200:1, v/v/v). After adding 20 μL of 5 M KCl, phase separation was made by adding 70 μL of chloroform. After spotting the lower chloroform phase onto the TLC plate, the TLC plate was developed with 1-butanol/ethanol/glacial acetic acid/H_2_O (8:2:1:2) in a glass tank. The S1P spot, which has a relative migration (R_f_) of approximately 0.72, was quantified after adding 600 μL of chloroform/methanol/water (5:5:1) using a fluorophotometer (excitation 460 nM, emission 534 nM). Sphingosine kinase activity was calculated based on the S1P spot intensity. All samples were prepared in triplicate, and the assay was repeated at least three times.

### Measurement of ROS

Superoxide anion level was determined using NBT solution (0.2%) in 50 mM sodium phosphate buffer (pH 7.5), and H_2_O_2_ level was determined using DAB staining solution (1 mg/ml) in distilled water. For total ROS determination, leaf epidermal strips were peeled from tobacco leaves and were floated on a solution of 50 μM DCFH-DA (Sigma Chemicals, St Louis, MO, USA). The leaf stripe samples were also co-treated with a solution consisting of 1 μM PHS. ROS was observed by fluorescence microscopy (excitation: 450 ± 490 nm; barrier 520 ± 560 nm) equipped with a cooled CCD camera (OLYMPUS, FV300, Japan).

### Ethylene Measurement

Ethylene production by PHS-treated plants was measured by gas chromatography (Hewlett Packard 5890 Series II, Wilmington, DE, USA) using an activated alumina column at 250°C and a flame ionization detector.

### Trypan Blue Staining

To monitor plant cell death, tobacco leaves were stained as previously described (Seo et al., 2020). PHS-treated tobacco whole leaves were immersed for 1 min in a boiling solution of 10 mL of lactic acid, 10 mL of glycerol, 10 g of phenol, and 0.4% (w/v) trypan blue. Stained plants were decolorized overnight and then photographed using a digital camera.

### Quantitation of Sporangia

Pathogen isolation from *Ppn*-infected tobacco leaves was performed following a modified method from that in a previously reported protocol (Sharma and Ghosh, 2016). Pathogen-infected tobacco leaves were was disinfected using a 2% NaClO (v/v) solution. After sterilized leaves were flooded with phosphate-buffered saline (PBS) at 4°C for 30 min and then incubated at 25°C for 16–20 h. After sonication, the number of sporangia was determined by using a hemocytometer.

## ACKNOWLEDGMENTS

This work was supported by grants from the National Research Foundation of Korea (Project No. NRF-2017R1D1A3B03034134) and Korean Research Institute of Biology and Biotechnology (Project No. 2019-0240) to K.Y.P.

## Supplemental data

The following material is available in the online version of this article.

**Table S1**. Primers used in this study for real-time qRT-PCR.

**Figure S1.** Effect of PHS on ROS accumulation in guard cells of tobacco leaves.

**Figure S2**. Effect of PHS on ROS accumulation in guard cells of *Ppn*-inoculated tobacco leaves.

**Figure S3**. Effect of PHS on expression profiles of *ACS* isoforms in *Ppn*-inoculated tobacco leaves.

**Figure S4**. Kinetics of *SphK* transcription in response to virulent *Ppn* attack and elicitor cryptogein treatment.

**Figure S5**. Transcription levels of *SphK* in *Ppn*-infected WT and transgenic plants after treatment with PHS.

## Supplementary Figure Legend

**Figure S1.** Effect of PHS on ROS accumulation in guard cells of tobacco leaves. A, ROS accumulation in guard cells was determined using confocal laser scanning microscopy (CLSM) after staining with 50 μM DCFH-DA. The CLSM images of DCF fluorescence (green) were merged with the bright-field images in the third column. Scale bars = 20 μm. B, Quantitation of DCF fluorescence intensity was calculated using ImageJ software.

**Figure S2**. Effect of PHS on ROS accumulation in guard cells of *Ppn*-inoculated tobacco leaves. The CLSM images of DCF fluorescence (green; DCFDA staining) and chlorophyll (Chl) autofluorescence (red) were determined at the indicated time after PHS treatment under *Ppn* inoculation. Both CLSM images were merged in the third column while the images in the bright field were merged in the fourth column. White boxes denote nuclei. Images are representative of three independent experiments with more than ten CLSM images at each indicated time. Scale Bars = 20 μm.

**Figure S3**. Effect of PHS on expression profiles of *ACS* isoforms in *Ppn*-inoculated tobacco leaves. Transcription levels of *NtACS* gene family members *NtACS1* (left panel), *NtACS2* (middle panel), and *NtACS4* (right panel) at the indicated times after treatment with 1 μM PHS in *Ppn*-infected tobacco leaves. Transcription levels are expressed relative to the reference gene *β-actin* after qRT-PCR. Ethylene levels and relative mRNA expression levels are expressed as means ± SD. An asterisk indicates a significant difference in PHS-treated cases or PHS and *Ppn* inoculation co-treated cases from mock-treated cases (one asterisk (P < 0.05) or two asterisks at the same time point (P < 0.01)).

**Figure S4.** Kinetics of *SphK* transcription in response to virulent *Ppn* attack and elicitor cryptogein treatment. Transcription levels of the *SphK* gene in tobacco plants after treatment with pathogen or cryptogein. Truncated tobacco plants with several leaves were shoot-inoculated with *Ppn*. Whole leaves of WT plants were treated with cryptogein (20 nM) for 72 h. Transcription levels are expressed relative to the reference gene *β-actin* after qRT-PCR. Relative mRNA expression levels are expressed as means ± SD. Asterisks indicate a significant difference in PHS-treated or co-treated cases with PHS and *Ppn* infection from mock-treated cases at the same time point (two asterisks (P < 0.01)).

**Figure S5.** Transcription levels of *SphK* in *Ppn*-infected WT and transgenic plants after treatment with PHS. The effects of PHS on transcription levels of *SphK* were determined in wild-type (WT) and transgenic plants with impaired ethylene biosynthesis (*CAS-AS*) and ROS production (*RbohD-AS* and *RbohF-AS*) after shoot inoculation with *Ppn* for 1 h or 48 h. Transcript amounts are expressed relative to the reference gene *β-actin* after qRT-PCR. Relative mRNA expression levels are expressed as means ± SD. Asterisks indicate a significant difference in PHS-treated or co-treated cases with PHS and *Ppn* infection from mock-treated cases at the same time point (one asterisk (P < 0.05) or two asterisks (P < 0.01)).

